# Frontoparietal theta synchronization causally links working memory with impulsive decision making

**DOI:** 10.1101/2024.10.17.618861

**Authors:** Georgia E. Kapetaniou, Gizem Vural, Alexander Soutschek

## Abstract

Delaying gratification in value-based decision making is canonically related to activation in the dorsolateral prefrontal cortex (dlPFC), but past research neglected that the dlPFC is part of a larger frontoparietal network. It is therefore unknown whether the dlPFC causally implements delay of gratification in concert with posterior parts of the frontoparietal network rather than in isolation. Here, we addressed this gap by testing the effects of frontoparietal theta synchronization and desynchronization on impulsive decision making using transcranial alternating current stimulation (tACS). Healthy participants performed an intertemporal choice task and a 3-back working memory task while left frontal and parietal cortices were stimulated with a 5 Hz theta frequency at in-phase (synchronization), anti-phase (desynchronization), or sham tACS. We found frontoparietal theta coupling to improve working memory performance, while in the decision task desynchronization was associated with more impulsive choices and stronger hyperbolic discounting of future rewards. Overall, our findings overcome the past focus of the dlPFC in isolation and show that patient decision making causally relies on synchronous activation in a frontoparietal network related to working memory.

## Introduction

The ability to delay gratification is a hallmark of individual success and psychological health (Baumeister, 2002; Warren K Bickel et al., 2019; Daugherty & Brase, 2010). A large body of evidence ascribes the dorsolateral prefrontal cortex (dlPFC) a central role for resisting immediate rewards in order to achieve long-term goals (Figner et al., 2010; McClure, Laibson, Loewenstein, & Cohen, 2004; Wesley & Bickel, 2014). However, as the dlPFC is part of the frontoparietal control network (Domenech, Redouté, Koechlin, & Dreher, 2018; Vincent, Kahn, Snyder, Raichle, & Buckner, 2008), it seems implausible to assume that dlPFC implements patience in intertemporal decisions in isolation rather than in concert with posterior regions like posterior parietal cortex (PPC). However, the majority of previous neural studies, and brain stimulation research in particular (Yang, Mauer, Vollm, & Khalifa, 2020; Yang, Vollm, & Khalifa, 2018), focused on the dlPFC either in isolation or in interaction with the neural reward system. Therefore, it remains unknown whether delaying gratification causally requires the dlPFC to synchronize activity with PPC.

Synchronous firing in dlPFC and PPC was already shown to play an important role for working memory processes, which in turn are hypothesized to contribute to delay of gratification (Hofmann, Schmeichel, & Baddeley, 2012). Synchronous, in-phase stimulation of dlPFC and PPC with transcranial alternating current stimulation (tACS) in the theta band enhanced working memory functioning, while desynchronous, out-of-phase stimulation impaired it (Alekseichuk, Pabel, Antal, & Paulus, 2017; Polania, Nitsche, Korman, Batsikadze, & Paulus, 2012; Violante et al., 2017). This suggests a causal role of functional coupling between dlPFC and PPC for working memory processes. Importantly, working memory has been linked to patience in intertemporal choice, as working memory processes may allow maintaining abstract information like the value of long-term rewards in mind during intertemporal decisions. Behaviorally, working memory capacity predicts patience in intertemporal choice (Hofmann et al., 2012), and neurally the brain correlates of working memory and intertemporal choice were found to strongly overlap (Jimura, Chushak, Westbrook, & Braver, 2018; Wesley & Bickel, 2014). We therefore hypothesized that strengthening control processes via synchronization versus desynchronization of frontoparietal activity promotes choices of long-term rewards.

To test our hypotheses, we synchronized and desynchronized theta band oscillations in dlPFC and PPC with tACS while participants performed a working memory and an intertemporal decision task. We expected to replicate previous findings that frontoparietal synchronization, relative to desynchronization, enhances working memory performance. We furthermore predicted that synchronization versus desynchronization of frontoparietal theta oscillations increases preferences for delayed over immediate rewards in the intertemporal decision task. This would provide evidence for a causal involvement of frontoparietal synchronization in delay of gratification and neurally link patient intertemporal decisions to working memory functioning.

## Methods

### Participants

30 healthy volunteers (mean age = 24 years, sd = 2.88, 15 female, 15 male) were recruited through the participant pool of the Munich Experimental Laboratory for Economic and Social Sciences (MELESSA) at the Ludwig Maximilian University Munich, Germany. The sample size was determined with an apriori power analysis (power = 80%, alpha = 5%) assuming an effect size of Cohen’s d = 0.47 from a previous study investigating the effect of phase synchronization on working memory (Alekseichuk et al., 2017). Volunteers were screened for counterindications to tACS prior to participation. Written informed consent was provided by all volunteers before the start of the experiment. Participants received a fixed compensation of 20 euros and an additional bonus depending on their decisions in the decision making task (see below). The study was approved by the local ethics committee of the psychology department at the University of Munich.

### Stimuli and task design

#### Working memory task

Participants performed a 3-back working memory task. During the task, participants viewed a sequence of letters, and each letter was presented on the screen for 1.5 seconds (inter-stimulus interval: 1.5 s; Figure 1A). The task was to indicate whether the letter currently displayed on the screen was identical with the letter presented 3 trials before (target stimulus). Participants were instructed to press the space button only if the current letter was a target.

**Figure 1.**
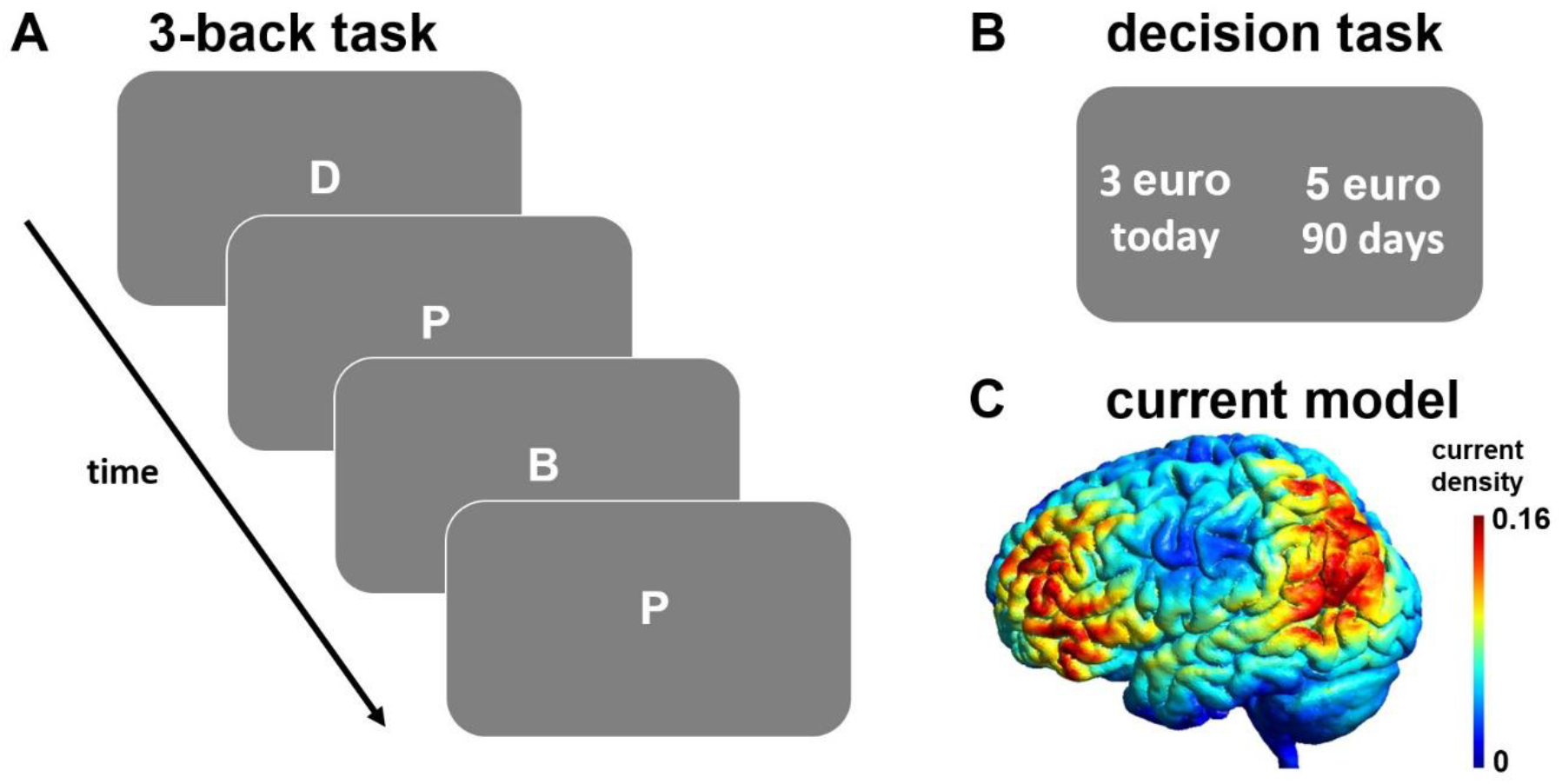
Experimental procedures. (A) In the 3-back working memory task, participants had to decide whether the currently presented letter is identical with the letter presented three trials before, requiring them to constantly maintain and update information in their working memory. (B) In the intertemporal decision task, participants made choices between smaller-sooner (e.g., 3 euro delivered today) and larger-later (e.g., 5 euro after 90 days) rewards. (C) Participants performed these tasks while undergoing in-phase (synchronizing), anti-phase (desynchronizing), or sham tACS over the left prefrontal and parietal cortex.

#### Intertemporal choice task

Participants performed an intertemporal choice task where they chose between two monetary rewards that were available at different points in time: a smaller-sooner (SS) reward, which was delivered at the end of the experiment, and a larger-later (LL) reward, which was delivered at a later date (Figure 1B. The SS reward ranged from 0.5 to 4.5 euro in steps of 0.5 euro, the LL was fixed at 5 euro and was delivered after 1 to 240 days. The two options were presented randomly on the left and right side of the screen and participants were asked to indicate their choice by pressing the left or right arrow key for the option on the left and right screen side, respectively, on a standard keyboard. Participants had 4 seconds to indicate their choice. After each decision, a fixation cross appeared on the screen for the remaining time of the 4 seconds, then the next trial started.

### Procedure

Participants performed the 3-back and the intertemporal choice task in three tACS conditions: frontoparietal synchronization, desynchronization, and sham (within-subject design). The intertemporal choice task included a total of 180 trials (60 trials per tACS condition), and the working memory task a total of 240 trials (80 per tACS condition). The tasks were administered in counterbalanced order and were performed in 12 miniblocks (2 per task and tACS condition). At the start of each miniblock, participants were stimulated with tACS for 30 seconds (current ramp-up: 5 seconds) without performing the task, followed by 120 seconds of task performance under stimulation. At the end of each miniblock, participants indicated whether they experienced any discomfort or flickering sensations due to the stimulation on a rating scale ranging from 0 (not at all) to 10 (very strongly). In the sham condition, the current was ramped down prior to task performance. Participants had a 30 second task- and stimulation-free break between the miniblocks to minimize potential carry-over effects between stimulation blocks (Christian, Kapetaniou, & Soutschek, 2023; Moisa, Polania, Grueschow, & Ruff, 2016; Soutschek, Moisa, Ruff, & Tobler, 2021).

At the end of the experiment, participants filled in demographic questionnaires and were debriefed. For the payment, one trial from the intertemporal choice task was randomly selected and implemented: if the participant had chosen the SS option in that trial, the corresponding amount was added as bonus to the standard compensation, whereas if they chose the LL option 5 euro were sent to them on the corresponding date via mail.

### tACS protocol

We applied tACS using a 4-channel tDCS stimulator (DC-Stimulator MC, neuroConn, Ilmenau, Germany). As in Biel, Sterner, Roll, and Sauseng (2022), we employed a high-definition 2×1 electrode set up. For the dlPFC we placed the active electrode over position F3 and the reference electrodes over positions Fz and F7 according to the international 10-20 system. For the PPC, the active electrode was placed over electrode position P3 and the reference electrodes over electrode positions Pz and P7. We used square rubber electrodes (3×3cm), which were attached to the participants’ head with the Ten20 conductive paste (Ten20 EEG Conductive Paste, Weaver and Company) and were kept steady throughout the session using fixation bandages. We performed current modeling using the Simnibs 2.1 toolbox (Saturnino et al., 2019), which suggested that this electrode set up led to strong and focal electrical fields in dlPFC and PPC, while stimulation effects between the two areas were negligible (Figure 1C). We stimulated participants in three conditions: synchronization (in-phase theta band (5 Hz) stimulation of dlPFC and PPC), desynchronization (anti-phase theta band stimulation of the two areas), and sham with a current strength of 1.5 mA peak-to-peak.

### Statistical analysis

In the *3-back task*, we analyzed tACS effects on reaction times and the sensitivity index d’ (difference between z-transformed hits and false alarms: Z_hits_ minus Z_false alarms_ (Soutschek & Tobler, 2020; Westbrook, Kester, & Braver, 2013)) with paired-samples *t*-tests.

In the *intertemporal choice task*, we performed both model-free and model-based analyses. For the model-free analysis, we used generalized linear mixed models (GLMMs) implemented via the lme4 package in R (Bates, Mächler, Bolker, & Walker, 2014) to regress binary choices (0 = SS option, 1 = LL option) on predictors for tACS (synchronization versus sham and desynchronization versus), amount of SS reward, and delay of LL reward. All continuous variables were z-standardized, and all fixed-effect predictors were also modelled as random slopes in addition to participant-specific random intercepts. As control variables of no interest, we added random slopes for stimulation-induced discomfort and flickering.

For the model-based analysis, we fitted hyperbolic discount functions to the choice data in a hierarchical Bayesian fashion using the JAGS software package (Plummer, 2003). We assumed that the subjective value of delayed rewards can be described by a canonical hyperbolic discount function (Frederick, Loewenstein, & O’Donoghue, 2002; Laibson, 1997):

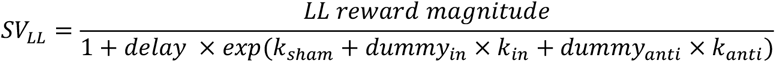

Where k_sham_ represents the hyperbolic discount factor (log-transformed to facilitate parameter estimation) under sham, whereas k_in_ and k_anti_ indicate the shift in hyperbolic discounting under synchronization (in-phase) and desynchronization (anti-phase), respectively, compared to sham. Subjective values were fitted to binary choices with a softmax link function including an inverse temperature parameter β as measure of choice consistency:

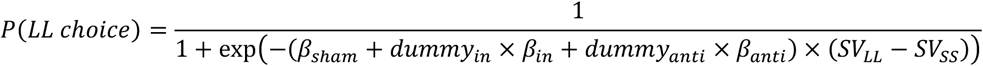

We fitted parameters both on the group and the individual level by assuming that individual parameters are normally distributed around the group means. To estimate the models, we used non-informative uniform priors and two chains with 25,000 iterations (10,000 burn-in samples). For all group parameters 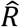 was ≤1.01, indicating model convergence. For statistical inference, we checked whether 95% highest-density interval (HDI) of the group-level posterior distributions included zero.

## Results

### Frontoparietal theta synchronization improves working memory performance

As manipulation check, we first assessed whether frontoparietal stimulation affected working memory performance. Sensitivity d’ as measure of performance accuracy in the n-back task (hits minus false alarms) was significantly increased under synchronization relative to sham, *t*(29) = 3.59, *p* = 0.001, Cohen’s d = 0.66, and desynchronization, *t*(29) = 3.59, *p* = 0.001, Cohen’s d = 0.66, whereas desynchronization showed no significant difference to sham, *t*(29) = 0.08, *p* = 0.93, Cohen’s d = 0.01 (Figure 2A). Thus, our results replicate previous findings on the causal involvement of frontoparietal synchronization in working memory (Alekseichuk et al., 2017; Polania et al., 2012).

**Figure 2.**
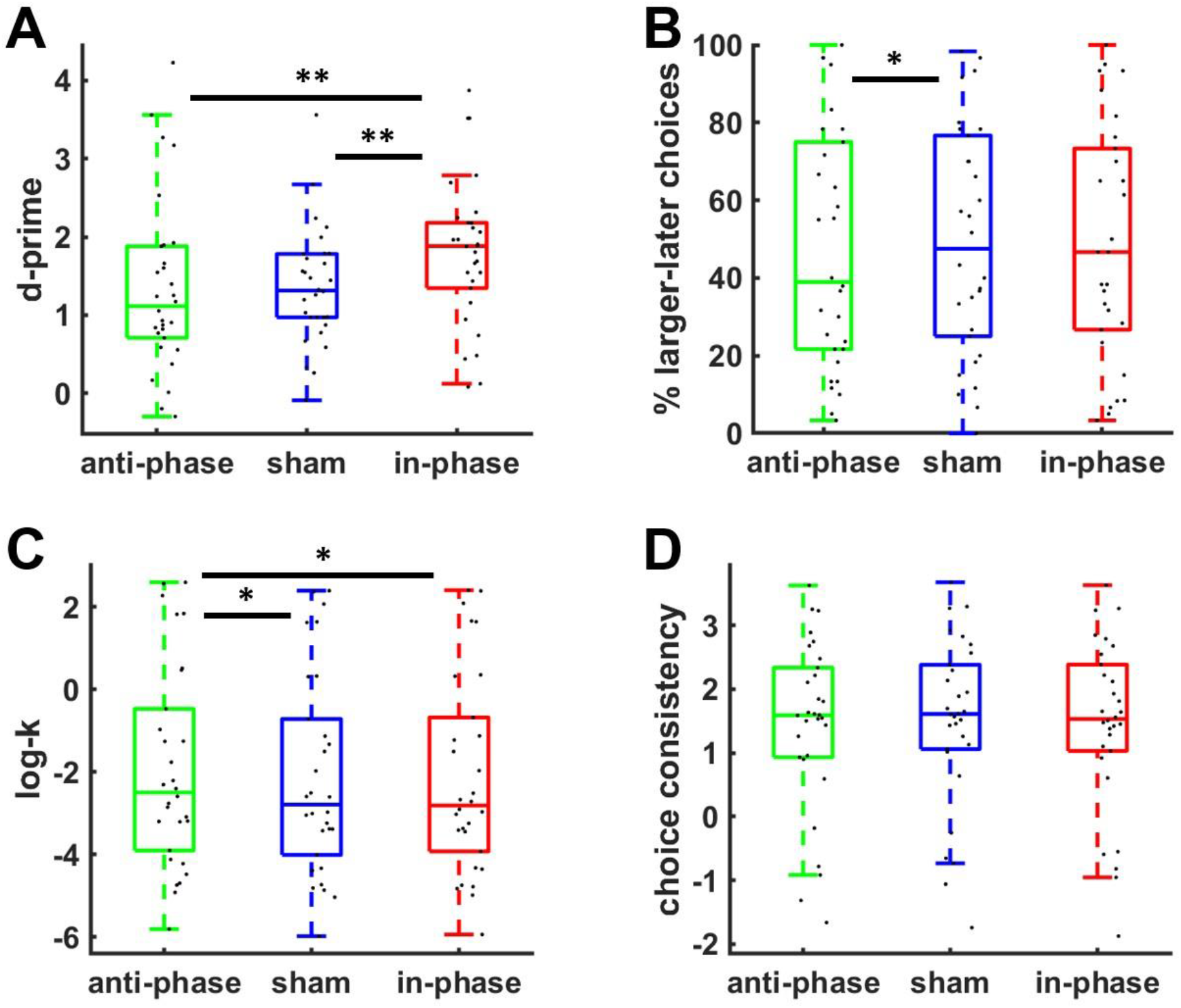
(A) Frontoparietal in-phase theta stimulation significantly improved working memory performance (sensitivity d’) compared to sham and anti-phase tACS. In the intertemporal decision task, anti-phase tACS resulted in (B) less choices of larger-later rewards and (C) stronger temporal discounting compared with control tACS. (D) We observed no stimulation effects on choice consistency (inverse temperature). For illustration purpose, (C) and (D) show extracted individual parameter estimates from the hierarchical Bayesian model. Black dots indicate individual data points; asterisks indicate significant effects (* *p* < 0.05; ** *p* < 0.01).

### Frontoparietal theta desynchronization decreases patient decision making

Based on the hypothesized link between working memory and the ability to delay gratification, we next assessed stimulation effects on intertemporal decisions. A model-free GLMM revealed that – as to be expected – the probability of choosing the LL option decreased with increasing amounts of the SS reward, beta = -2.16, *z* = 10.52, *p* < 0.001, and with longer delays until LL reward delivery, beta = -1.85, *z* = 7.30, *p* < 0.001. While we observed no influence of synchronization relative to sham on choices, beta = 0.02, *z* = 0.12, *p* = 0.91, theta desynchronization significantly reduced preferences for delayed rewards compared with sham, beta = -0.30, *z* = 2.04, *p* = 0.04, though not compared with synchronization, beta = 0.09, *z* = 0.44, *p* = 0.66 (Figure 2B). This supports the hypothesized involvement of frontopolar theta coupling in intertemporal decision making.

The model-free results are corroborated by a model-based analysis of hierarchically estimated hyperbolic discount factors (Table 1). Desynchronization significantly increased hyperbolic discounting of delayed rewards, HDI_mean_ = 0.20, HDI_95%_ = [0.03, 0.36], while synchronization showed no significant effects, HDI_mean_ = 0.01, HDI_95%_ = [-0.14, 0.17]. A direct comparison between synchronization and desynchronization suggested that desynchronization increased delay discounting also relative to theta synchronization, HDI_mean_ = 0.18, HDI_95%_ = [0.01, 0.34] (Figure 2C). There was no evidence for stimulation effects on choice consistency, as the inverse temperature parameter was unaffected by either synchronization, HDI_mean_ = -0.05, HDI_95%_ = [-0.23, 0.15], or desynchronization, HDI_mean_ = -0.05, HDI_95%_ = [-0.24, 0.15] (Figure 2D). Together, this provides converging evidence that frontoparietal theta desynchronization increases impulsiveness in intertemporal choice.

**Table 1.** Results for the hierarchically estimated hyperbolic discount model. The parameters k_sham_, k_in_, k_anti_ refer to the posterior distributions of the group-level log-transformed hyperbolic discount factors under sham, synchronization versus sham, and desynchronization versus sham, respectively, whereas β_sham_, β_in_ and β_anti_ refer to the inverse temperature (choice consistency) parameter. Standard errors of the mean are in brackets.

| Parameter | Mean | HDI<sub>2.5%</sub> | HDI<sub>97.5%</sub> |
| --- | --- | --- | --- |
| $k_{\text{sham}}$ | -1.92 (0.55) | -2.97 | -0.79 |
| $k_{\text{in}}$ | 0.01 (0.08) | -0.14 | 0.17 |
| $k_{\text{anti}}$ | 0.20 (0.09) | 0.03 | 0.36 |
| $\beta_{\text{sham}}$ | 1.51 (0.28) | 0.97 | 2.04 |
| $\beta_{\text{in}}$ | -0.05 (0.09) | -0.23 | 0.15 |
| $\beta_{\text{anti}}$ | -0.05 (0.10) | -0.24 | 0.15 |

Finally, based on studies suggesting a link between working memory and intertemporal decisions, we tested for the hypothesized correlation between working memory performance (sensitivity d’) and delay discounting (individual parameter estimates from hyperbolic discount model). Under sham, better working memory performance (d’) was associated with weaker temporal discounting (log-k), Spearman’s rho = -0.39, *p* = 0.02, one-tailed (Figure 3A). Moreover, the baseline-corrected influences of synchronization versus desynchronization on d’ and log-k were correlated, Spearman’s rho = -0.34, *p* = 0.03, one-tailed (Figure 3B): individuals with stronger working memory improvement under synchronization versus desynchronization showed also more patient decisions (more negative log-ks) under synchronization versus desynchronization. This suggests a possible link between the tACS effects on working memory and decision making.

**Figure 3.**
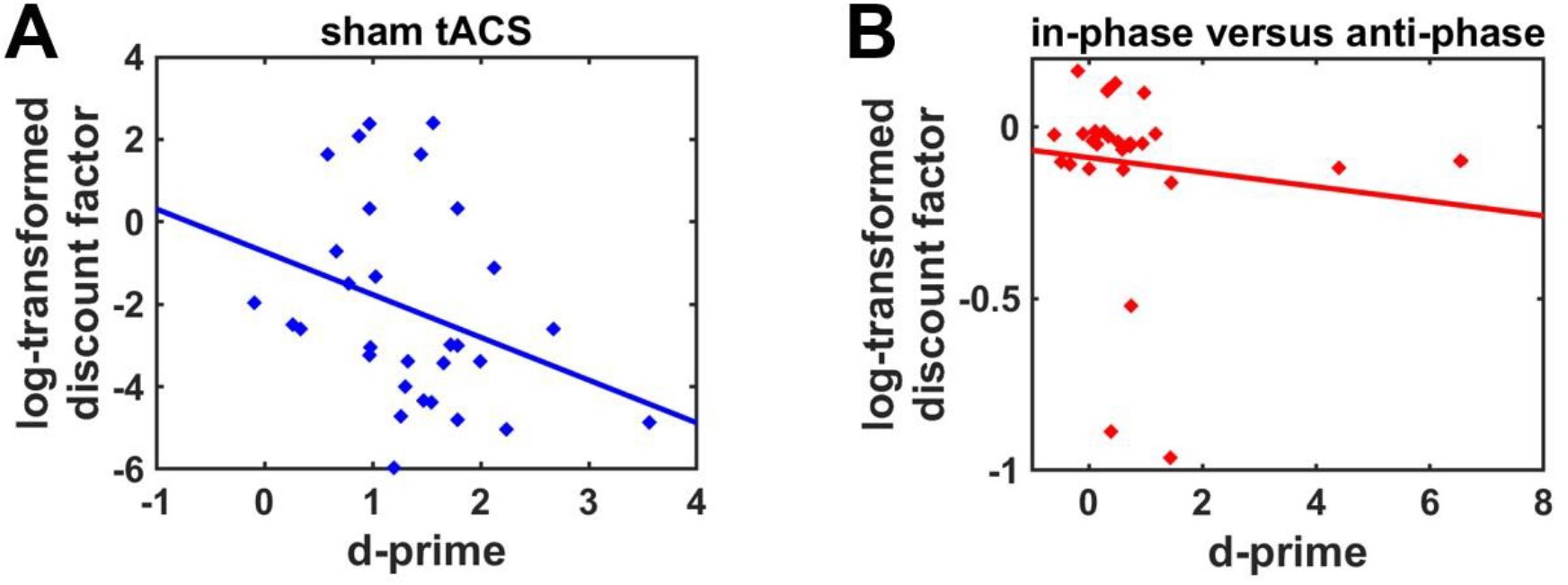
Correlations between working memory performance (sensitivity d’) and log-transformed discount parameters (A) under sham and (B) under synchronization versus desynchronization (baseline-corrected). Note that statistical inferences are based on non-parametric tests to account for the skewedness of the parameter distributions.

## Discussion

Frontoparietal theta coupling plays an important role in working memory functioning, but little is known about its contribution to value-based decision making. Here, we show that desynchronization of frontoparietal theta oscillations enhances the discounting of delayed rewards, suggesting a causal role of frontoparietal coupling for intertemporal choice. Because synchronization versus desynchronization of frontoparietal oscillations also improved working memory performance, replicating previous findings (Alekseichuk et al., 2017; Biel et al., 2022; Polania et al., 2012), our findings suggest overlapping neural mechanisms to underlie working memory and patient decision making. This is further evidenced by significant correlations between working memory performance and delay discounting. Taken together, our results suggest frontoparietal theta coupling to causally underlie both working memory and decision processes.

By highlighting the importance of synchronized brain network activity, our findings go beyond current neural models of intertemporal decision making: Previous research ascribed the dlPFC a role for encoding long-term goals and for modulating the subjective value of rewards in the brain’s reward system (Hare, Hakimi, & Rangel, 2014; Smith, Monterosso, Wakslak, Bechara, & Read, 2018; van den Bos, Rodriguez, Schweitzer, & McClure, 2014; Wesley & Bickel, 2014). However, this perspective neglected that the dlPFC is part of a frontoparietal control network (Vincent et al., 2008), making it reasonable to assume that the dlPFC implements patient decisions not in isolation but in interaction with the parietal cortex. In fact, past studies provided evidence for PPC activation during intertemporal decisions (Boettiger et al., 2007; Rodriguez, Turner, Van Zandt, & McClure, 2015) or also more ventral parts of the parietal cortex (Soutschek, Moisa, Ruff, & Tobler, 2020; Soutschek, Ruff, Strombach, Kalenscher, & Tobler, 2016), but evidence for dlPFC-PPC connectivity during decision making was lacking so far. Our findings fill this gap by showing that the dlPFC’s influence on intertemporal choice requires synchronization with lateral PPC. Note that one previous study provided evidence for a causal involvement of medial (rather than lateral) frontoparietal synchronization for value-based decision making, positing that the network performs value-to-action transformations communicated from the PFC to the PPC to translate values into actions (Polania, Moisa, Opitz, Grueschow, & Ruff, 2015). We assume that the lateral network identified in the current study may play a similar role in intertemporal choice: dlPFC-PPC synchronization may promote the transfer of information about delayed reward values encoded in dlPFC to the PPC, where the values are assigned to action options (Sugrue, Corrado, & Newsome, 2004). As caveat, we note that the current findings provide no insights into the directionality of the information flow between dlPFC and PPC. Nevertheless, our results highlight that the dlPFC contributes to value-based choice as part of a frontoparietal network, going beyond prevalent views in the literature (Smith et al., 2018; Wesley & Bickel, 2014; Yang et al., 2018).

The influence of frontoparietal coupling on decision making moreover appears to be related to working memory processes. Consistent with past research (Hofmann et al., 2012), better working memory performance was associated with less impulsive decision making, and the stimulation effects on delay discounting co-varied with tACS-induced working memory improvements. Delaying gratification is thought to require the representation of the value of long-term rewards (which are less concrete than immediately available outcomes (Fujita, 2011; Stillman et al., 2017)) in working memory (Jimura et al., 2018; Smith et al., 2018; Wesley & Bickel, 2014). Our findings provide neural support for theoretical accounts on the role of working memory for delay of gratification by suggesting an involvement of working memory-related frontoparietal coupling in intertemporal decision making.

Taken together, our results show that working memory-related frontoparietal theta coupling causally implements patience in impulsive decision making. This provides a network perspective on the contribution of the neural control system to decision making, overcoming the focus of past research on the DLPFC in isolation (or in interaction with the subcortical reward system). Given the prevalence of impulsive decision making in several clinical disorders (W. K. Bickel, Koffarnus, Moody, & Wilson, 2014; Monterosso, Piray, & Luo, 2012; Stutzer & Meier, 2015; Volkow & Baler, 2015), these findings may contribute to the development of more effective neural treatments of impulsiveness.

## Acknowledgements

We kindly thank the Munich Experimental Laboratory for Economic and Social Sciences (MELESSA) for support with recruitment and data collection.

## Data availability statement

The raw data underlying the reported reports will be publically available on the Open Science Framework (OSF).

